# Discovery of novel papillomaviruses in the critically-endangered Malayan and Chinese pangolins

**DOI:** 10.1101/2022.10.04.510846

**Authors:** Jose Gabriel Nino Barreat, Anselmo Jiro Kamada, Charles Reuben de Souza, Aris Katzourakis

## Abstract

Pangolins are scaly and toothless mammals which are distributed across Africa and Asia. Currently, the Malayan, Chinese and Philippine pangolins are all designated as critically-endangered species. Although few pangolin viruses have been described, their viromes have received more attention following the discovery that they harbour sarbecoviruses related to SARS-CoV-2. Using a large-scale genome mining strategy, we discovered novel lineages of papillomaviruses infecting the Malayan and Chinese pangolins. We were able to assemble 3 complete circular papillomavirus genomes with an intact coding capacity, and 5 additional L1 genes encoding the major capsid protein. Phylogenetic analysis revealed that 7 out of 8 L1 sequences formed a monophyletic group which is the sister lineage to the Tree shrew papillomavirus 1, isolated from Yunnan province in China. Additionally, a single L1 sequence assembled from a Chinese pangolin was placed in a clade closer to alpha- and omegapapillomaviruses. Examination of the SRA data from 95 re-sequenced genomes revealed that 49.3% Malayan pangolins and 50% Chinese pangolins, were positive for papillomavirus reads. Our results indicate that pangolins in South East Asia are the hosts to diverse and highly prevalent papillomaviruses, which may have implications for pangolin health and conservation.

## Introduction

Pangolins are scaly, primarily nocturnal and insectivorous mammals which belong to the order Pholidota. They are classified into three genera, two of which are found in Africa (*Smutsia*, *Phataginus*), and one in Asia (*Manis*) (1). There are four different species of pangolins in Asia: the Chinese pangolin (*Manis pentadactyla*), the Malayan pangolin (*Manis javanica*), the Indian pangolin (*Manis crassicaudata*) and the Philippine pangolin (*Manis culionensis*) (1). Of these, the Chinese, Malayan and Philippine pangolins are currently designated as critically-endangered species by the International Union for the Conservation of Nature (IUCN), due to their population decline as a result of overexploitation and trafficking for their scales (2–4).

Pangolins have recently gained attention as a potential host of emerging viral diseases after coronaviruses related to SARS-CoV-2 were reported in the Malayan pangolin (5–10). Pangolins have also been recognised as hosts for several other RNA viruses such as Canine distemper virus (*Paramyxoviridae*), which is associated with respiratory, digestive and neurological illness in pangolins (11), flaviviruses, reoviruses, pneumoviruses and picornaviruses (12–15). Overall, the diversity of DNA viruses and their disease association in pangolins is less well known; so far anellovirus, parvovirus, circovirus and genomovirus genomes have been described (15,16).

Papillomaviruses are non-enveloped, dsDNA viruses with a circular genome, which can cause a diverse array of clinical manifestations in their vertebrate hosts, ranging from subclinical, to cutaneous and mucosal warts, and cancerous lesions (17). We describe the discovery of two novel lineages of papillomaviruses found by mining the genome data of the Malayan and Chinese pangolins. These findings constitute the first detailed record of papillomavirus infection in pangolins, and highlight the need for a systematic assessment of the diversity and biology of DNA viruses hosted by these animals.

## Methods

We made the initial discovery of the unidentified contig (NW_023450026.1) in pangolins by querying the RefSeq eukaryotic genomes database (ref_euk_rep_genomes) with 1,413 papillomavirus reference proteins obtained from the NCBI Virus Resource (June/2022) (18,19). Screening was performed using the tblastn algorithm (−task tblastn-fast) implemented by the ElasticBLAST (v0.2.6) method on the Google Cloud Platform (20,21). The search returned 1,017 hits with e-values < 1e-5 to an unplaced genomic scaffold (YNU_ManJav_2.0 scaffold_14136) from the RefSeq genome assembly of the Malayan pangolin.

The 7,307-bp contig was annotated using the PuMA pipeline (22). This sequence corresponded to a full papillomavirus genome encoding L1, L2, E1, E2, E6 and E7, in addition to two spliced products (E1^E4 and E8^E2). Given that the papillomavirus genome was intact, we screened the short-read data of the re-sequenced genomes of 72 Malayan pangolin and 22 Chinese pangolin individuals, also produced by the study of Hu et al. (23). We obtained the SRA experiment accession numbers from this study (BioProject IDs: PRJNA529540, PRJNA529512), and used a combination of blastn and tblastn on the NCBI web server (24) to find reads with significant similarity to papillomaviruses in these experiments.

We downloaded the short-read sequences of the SRA experiments with more than 100 significant matches, and tried to *de novo* assemble complete viral genomes or the L1 gene. First, fasta files were concatenated and the duplicate sequences were removed with the rmdup option of SeqKit (25). Then we used a custom Python 3 script to sort the sequences into forward, reverse and orphan reads, since they were sequenced as paired-end libraries. The reads in these files were assembled in SPAdes v3.15.4 (26), using the “–metaviral” and “– assembler-only” flags. Contigs of complete viral genomes were annotated using PuMA. Contigs of L1 genes were examined in ORFfinder to check for the presence of an intact L1 open reading frame (27). The molecular weight and isoelectric point of predicted protein products were estimated in the ExPASy web server (28). We calculated the coverage (depth) of our assembled sequences using Magic-BLAST (29), a tool used for mapping next-generation DNA or RNA reads to a reference sequence. Mapped reads were sorted and the coverage calculated using samtools (30).

To study the systematics of these viruses, we inferred a Bayesian phylogeny of the L1 proteins. We first collected a set of L1 sequences more closely related to the pangolin sequences in searches of the PaVE papillomavirus taxonomy tool (31). We selected proteins from the type species of papillomaviruses recognised by the ICTV in addition to the Tree shrew papillomavirus 1 (and Tree shrew papillomavirus 2) described in the study of Liu et al. (32). Sequences were aligned in MAFFT v7.490 using the accurate option (mafft-linsi) (33). The multiple sequence alignment was trimmed manually, resulting in a final alignment of 522-aa. The best substitution model for the alignment (LG+I+G4+F) was found in ModelTest-NG (34). We then inferred a Bayesian phylogeny in MrBayes version 3.2.7a (35) with an MCMC chain length of 1,000,000 generations (burn-in = 25%). Convergence was assessed by ensuring that the Average standard deviation of split frequencies < 0.01 and the Potential scale reduction factor for all parameters was ~1.

## Results

We found significant hits to papillomaviruses in 36 out of 73 (49.3%) samples examined for the Malayan pangolin, and 11 out of 22 (50%) samples from the Chinese pangolin (Table 1 and Table S1). All the samples with known geographic origin were annotated as coming from the Yunnan province of China (sampled between the years 2000-2005). The remaining 42 individuals were seized either in Yunnan (yrs. 2016-2017) or at the border between Yunnan and Myanmar (Sino-burmese border, yr. 2014). From the positive individuals, 10 out of 73 (13.7%) Malayan pangolins, and 1 out of 22 (4.6%) Chinese pangolins had more than 100 significant hits. We selected these for assembly of the complete genome/L1 gene. Three complete papillomavirus genomes were assembled *de novo*: two new papillomavirus genome assemblies (MJ55: depth = 36.71x, MJ23: depth = 10.22x) and the reassembled genome for the reference contig (MJ74: depth = 13.06x). We could also assemble the complete L1 gene for three other individuals (MJ18, MJ33, MP15). Interestingly, three different L1 gene contigs were assembled from a single Chinese pangolin individual, designated as MP15A, MP15B and MP15C.

**Table 1.**
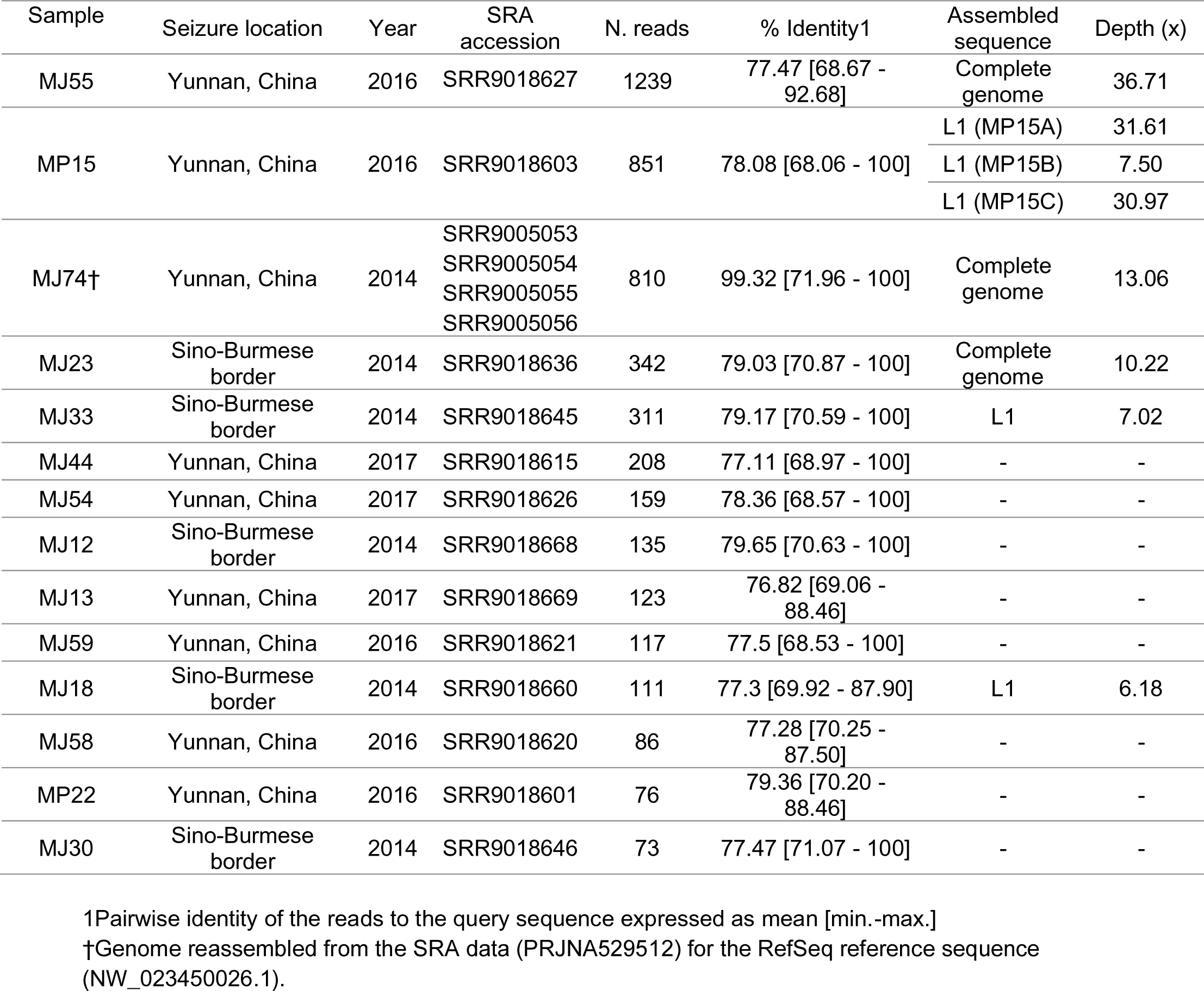
Summary of the SRA experiments from pangolins with >70 significant hits (blastn, e-value < 0.01) to the linearised papillomavirus genome of the reference individual MJ74. Sets of reads which could be assembled into a complete genome or complete L1 gene are indicated, with a reference to the mean depth (x) of the assembly. Samples with less than 70 positive reads are shown in Table S1.

The assembled genomes ranged in size from 7,253 bp to 7,437 bp with a GC content between 39.84 and 40.09%. All three genomes encode the four core papillomavirus proteins: E1, E2, L1 and L2. In addition, they encode the accessory proteins E6 and E7 (Figure 1, Figures S1 and S2). We were also able to identify two spliced products, E8^E2 and E1^E4, for the MJ74 and MJ23 papillomavirus genomes. The Upstream Regulatory Regions (URRs) range in size from 504 to 613 bp, and contain a binding site for the E1 and E2 proteins (E1BS 100% consensus: TTGTTGTTRGCAACAACMAT, E2BS 100% consensus: AACGAATTCGGTY). All the proteins are encoded in the forward strand as typical for papillomaviruses (17). A table with the gene coordinates, protein lengths, molecular weights and isoelectric points of predicted proteins is provided in the Supplementary materials (Table S2).

**Figure 1.**
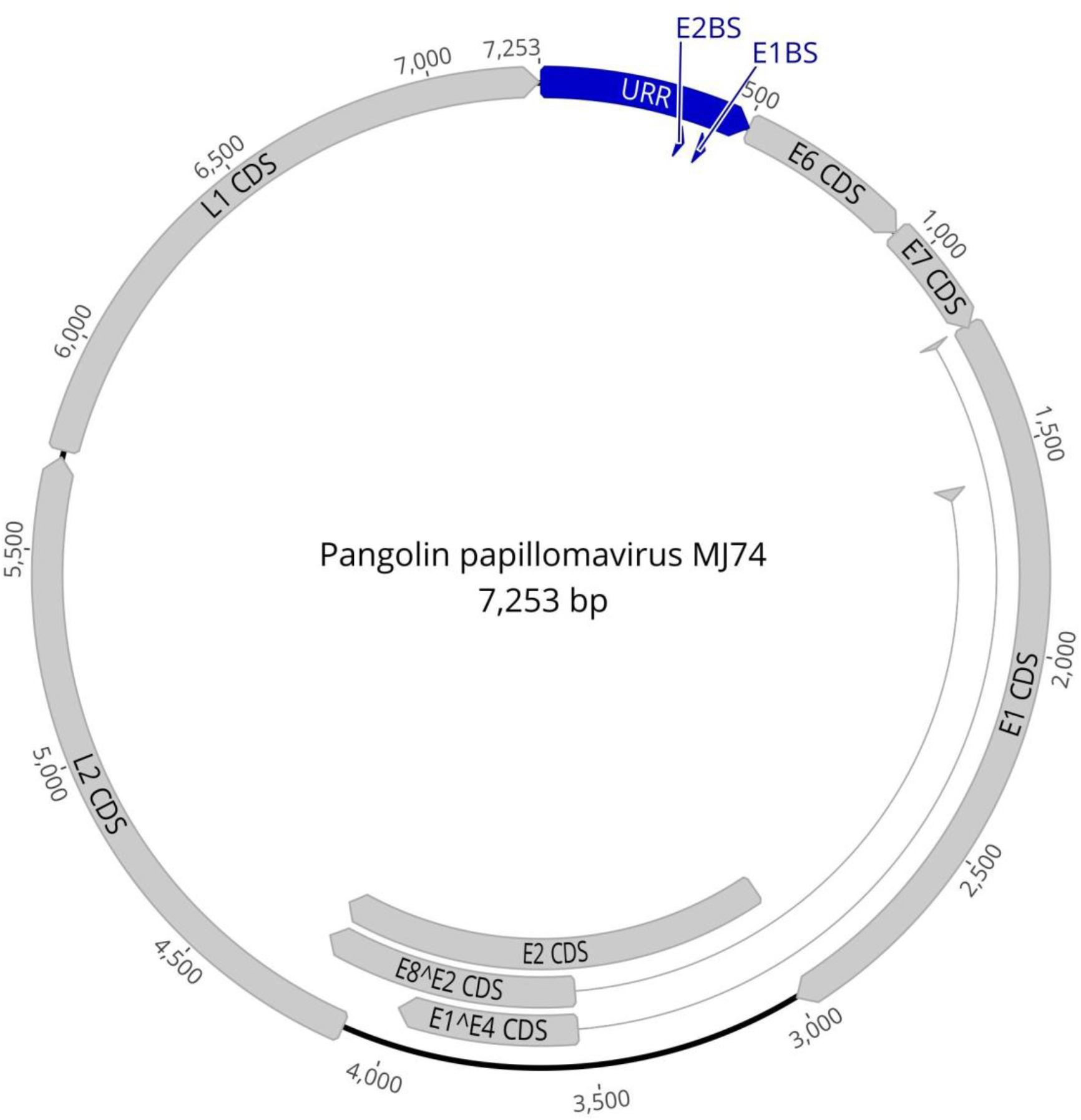
Complete papillomavirus genome assembled from the Malayan pangolin individual MJ74, and annotated with the PuMA pipeline. The 7,253-bp circular genome encodes the four core papillomavirus proteins (L1, L2, E1, E2) in addition to the E6 and E7 accessory proteins. Two spliced products were also identified: E1^E4 and E8^E2. The Upstream Regulatory Region (URR) has also been annotated and includes the E1- and E2-protein binding sites (E1BS, E2BS). All genome features and sequences are provided as a GenBank file (Supplementary information). Image created in Geneious Prime 2022.2.1 (36).

Phylogenetic analysis of the L1 proteins placed 7 out of 8 of the L1 sequences into a monophyletic group with high support (posterior probability, PP. = 1) (Figure 2). These sequences were grouped as the sister lineage to the L1 protein of the Tree shrew papillomavirus 1, also with high support (PP. = 1). The tree shrew and seven pangolin papillomavirus L1 sequences, were placed in the same clade as viruses in the genera *Kappapapillomavirus*, which infect both Old and New World rabbits, and *Dyodeltapapillomavirus* which infect North American beavers (PP. = 0.97). A single L1 protein sequence from a Chinese pangolin (MP15C), was nested in a different clade that includes the genera *Omegapapillomavirus* isolated from polar bears, *Alphapapillomavirus* which infects primates (including human papillomaviruses type 32/16/18) and *Dyodeltapapillomavirus* which infects domestic pigs (PP. = 1). However, the position of the sequence within the clade, basal to omega- and alphapapillomaviruses, had a low support (PP. = 0.61), so its specific placement within the clade is uncertain.

**Figure 2.**
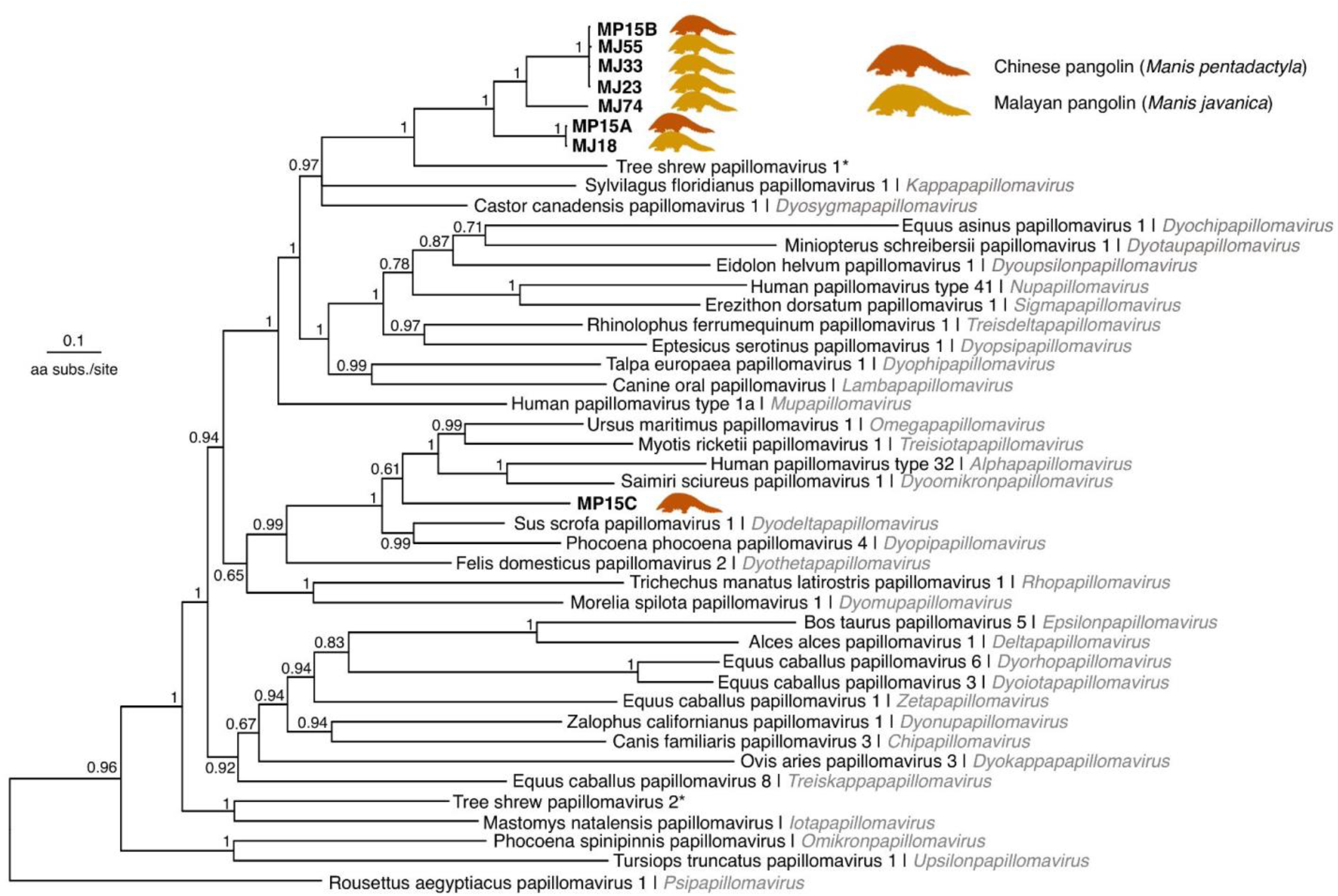
Bayesian phylogenetic tree based on the L1 proteins of papillomaviruses. Most pangolin papillomavirus L1 sequences form a highly supported monophyletic group which is the sister to the Tree shrew papillomavirus 1 (posterior probability = 1). This clade includes sequences assembled from 5 Malayan pangolin individuals (MJ18, MJ23, MJ33, MJ55, MJ74), and two sequences from a Chinese pangolin individual (MP15A, MP15B). A single sequence from the same Chinese pangolin individual (MP15C) was placed with high confidence (posterior probability = 1) on a different clade that includes alpha-, omega- and dyodeltapapillomaviruses. Tree estimated in MrBayes using the LG+I+G4+F amino acid substitution model, an MCMC chain length of 1,000,000 generations and a 25% burn-in. The tree was outgroup-rooted with the L1 sequence from Sparus aurata papillomavirus 1 (*Secondpapillomavirinae*), which is not shown for clarity.

## Discussion

We describe the complete genome for a novel papillomavirus, which constitutes the first detailed record of the family *Papillomaviridae* in pangolins. The assembly of complete circular genomes with an intact coding capacity, regulatory elements, and no host flanking sequences, shows that these are exogenous viruses highly likely to be infecting pangolins. In addition, finding these viruses across multiple individuals, sampled from different time-points and in two different host species, suggests the viruses naturally circulate in pangolin populations.

We found that 7 out of 8 of the predicted L1 protein sequences formed a highly supported clade, suggesting that this group belongs to a single species of papillomavirus present in the Chinese and Malayan pangolins (L1 nucleotide percent identities: 71.87-99.93%). This clade, which we call Pangolin papillomavirus 1, was placed as the sister group to Tree shrew papillomavirus 1; isolated from wild Northern tree shrews (*Tupaia belangeri*) in the Jianchuan and Lufeng localities of Yunnan province (yrs. 2016-2017) (32). Since the pangolin and Tree shrew papillomavirus 1 form a highly supported monophyletic group, share similar genome structures (both lack the protein E5), and infect mammals in the same geographic region (Figure S3), we consider they should be classified as members of the same genus. Even though traditional taxonomic criteria would place the Pangolin papillomavirus 1 and Tree shrew papillomavirus 1 in a single species (L1 nucleotide percent identities: 63.22%-65.81%, i.e. >60%) (37), we believe that from an evolutionary perspective it is valid to consider them as two separate species. In favour of this argument, the Pangolin papillomavirus 1 lineage forms a monophyletic group distinct from the Tree shrew papillomavirus 1 (Figure 2), and they infect distantly related hosts belonging to different orders of mammals (order Pholidota and order Scandentia). In addition, a time-calibrated phylogeny suggests this group of papillomaviruses has infected and adapted to their pangolin hosts for at least 526 (262-905) years, and diverged from the Tree shrew papillomavirus 1 lineage about 1001 (638-1674) years ago (Figure S4).

Noticeably, we assembled two different L1 sequences (MP15A, MP15B), belonging to the Pangolin papillomavirus 1 lineage from a single Chinese pangolin individual (MP15), along with another papillomavirus L1 sequence that clustered with omega- and alphapapillomaviruses clade instead (Figure 2, we refer to this virus provisionally as Pangolin papillomavirus 2). This suggests that the individual was likely co-infected by two distinct strains of Pangolin papillomavirus 1, in addition to the Pangolin papillomavirus 2. These results suggest that South East Asian pangolins may be potentially the hosts of papillomaviruses from diverse lineages, which will require further study and characterisation.

After examining the SRA samples for the re-sequenced genomes of 73 Malayan and 22 Chinese pangolins, we found positive reads for Pangolin papillomavirus 1 in about 50% of individuals in both species. The sequences were confirmed to be of pangolin origin since they were sampled from muscle tissue of pangolins (23) (Li Y., pers. communication), they were assembled to a good coverage, they show sequence variation, and have a unique phylogenetic placement in the papillomavirus phylogeny. Finding papillomavirus (SRA) positive pangolin individuals distributed across a timespan of 17 years indicates that papillomavirus infections are prevalent among these pangolin species.

We also found reads with significant matches (e-values: 9·10^−29^−2·10^−3^) to the Pangolin papillomavirus 1 genome in the samples from five individuals described in the meta-transcriptomic study of the Malayan pangolin by Shi et al. (15) (Table S3). The difference in the number of matching reads we obtained from the genomic and transcriptomic SRA samples may be attributed to latent papillomavirus infections and differences in sampling (different tissues were pooled together for the meta-transcriptomic work). We also examined an additional report of papillomavirus reads in the Malayan pangolin by Liu et al. (5). However, we did not obtain any matches to our query sequences, and found that the best-hits to their custom database were all human-HPV16 junctions (Table S4). No hits could be obtained using the reference HPV16 virus proteins in a tblastn, suggesting that the reads came from human DNA contamination.

We demonstrate the application of cloud-computing for the fast and efficient mining of large-scale genomic data sets, and the discovery of novel viral lineages. This data-driven virus discovery (DDVD) is expected to increase our knowledge of the virosphere and offer a springboard for the experimental characterisation of viruses that have not been isolated or which silently infect their hosts (38). Using these *in silico* screening methods, a new lineage of fish alloherpesvirus-like endogenous viruses has been discovered (39), in addition to millions of novel RNA viruses on a global scale (40–43). Given the critically-endangered conservation status of the Malayan and Chinese pangolins in South East Asia, it will be important to actively assess whether these viruses cause any disease or decrease the fitness of their pangolin hosts. It is also still unclear whether the Malayan pangolin populations in the islands of Borneo, Java and Sumatra, and the Philippine pangolin, also host similar papillomaviruses. Further studies into these questions may shed light on the impact papillomaviruses have on pangolins and inform potential strategies for conservation.

## Acknowledgements and funding

This work was funded by an ERC grant to A.K. (101001623-PALVIREVOL), and partly by a Google Research Grant to J.G.N.B.

## References

1. Gaubert P, Wible JR, Heighton SP, Gaudin TJ. Phylogeny and systematics. In: Challender D, Nash H, Waterman C, editors. Pangolins: Science, Society and Conservation. London: Academic Press: Elsevier; p. 630.

2. Challender D, Willcox DHA, Panjang E, Lim N, Nash H, Heinrich S, et al. Manis javanica. The IUCN Red List of Threatened Species 2019: e.T12763A123584856. [Internet]. Available from: https://dx.doi.org/10.2305/IUCN.UK.2019-3.RLTS.T12763A123584856.en

3. Challender D, Wu S, Kaspal P, Khatiwada A, Ghose A, Ching-Min S, et al. *Manis pentadactyla* (errata version published in 2020). The IUCN Red List of Threatened Species 2019: e.T12764A168392151. [Internet]. Available from: https://dx.doi.org/10.2305/IUCN.UK.2019-3.RLTS.T12764A168392151.en

4. Schoppe S, Katsis L, Lagrada L. Manis culionensis. The IUCN Red List of Threatened Species 2019: e.T136497A123586862. [Internet]. Available from: https://dx.doi.org/10.2305/IUCN.UK.2019-3.RLTS.T136497A123586862.en

5. Liu P, Chen W, Chen JP. Viral Metagenomics Revealed Sendai virus and coronavirus infection of Malayan pangolins (*Manis javanica*). Viruses. 2019 Nov;11(11):979.

6. Lam TTY, Jia N, Zhang YW, Shum MHH, Jiang JF, Zhu HC, et al. Identifying SARS-CoV-2-related coronaviruses in Malayan pangolins. Nature. 2020 Jul;583(7815):282–5.

7. Li L, Wang X, Hua Y, Liu P, Zhou J, Chen J, An F, Hou F, Huang W, Chen J. Front Microbiol. 2021 Mar 3;12:657439.

8. Yang S, Shan T, Xiao Y, Zhang H, Wang X, Shen Q, et al. Digging metagenomic data of pangolins revealed SARS-CoV-2 related viruses and other significant viruses. J Med Virol. 2021;93(3):1786–91.

9. Zhang T, Wu Q, Zhang Z. Probable pangolin origin of SARS-CoV-2 associated with the COVID-19 outbreak. Curr Biol. 2020 Apr 6;30(7):1346–1351.e2.

10. Xiao K, Zhai J, Feng Y, Zhou N, Zhang X, Zou JJ, et al. Isolation of SARS-CoV-2-related coronavirus from Malayan pangolins. Nature. 2020 Jul;583(7815):286–9.

11. Chin JSC, Tsao EH. Chapter 40: Pholidota. In: Miller ER, Fowler ME, editors. Fowler’s Zoo and Wild Animal Medicine. St. Louis, Missouri: Saunders; 2015. p. 369–75.

12. Yang R, Peng J, Zhai J, Xiao K, Zhang X, Li X, et al. Pathogenicity and transmissibility of a novel respirovirus isolated from a Malayan pangolin. J Gen Virol. 102(4):001586.

13. Gao WH, Lin XD, Chen YM, Xie CG, Tan ZZ, Zhou JJ, et al. Newly identified viral genomes in pangolins with fatal disease. Virus Evol. 2020 Jan 1;6(1):veaa020.

14. Wang X, Chen W, Xiang R, Li L, Chen J, Zhong R, et al. Complete genome sequence of Parainfluenza virus 5 (PIV5) from a Sunda pangolin (*Manis javanica*) in China. J Wildl Dis. 2019 Oct 1;55(4):947–50.

15. Shi W, Shi M, Que TC, Cui XM, Ye RZ, Xia LY, et al. Trafficked Malayan pangolins contain viral pathogens of humans. Nat Microbiol. 2022 Aug;7(8):1259–69.

16. Ning S, Dai Z, Zhao C, Feng Z, Jin K, Yang S, et al. Novel putative pathogenic viruses identified in pangolins by mining metagenomic data. J Med Virol. 2022;94(6):2500–9.

17. Van Doorslaer K, Chen Z, Bernard HU, Chan PKS, DeSalle R, Dillner J, et al. ICTV Virus Taxonomy Profile: *Papillomaviridae*. J Gen Virol. 99(8):989–90.

18. O’Leary NA, Wright MW, Brister JR, Ciufo S, Haddad D, McVeigh R, et al. Reference sequence (RefSeq) database at NCBI: current status, taxonomic expansion, and functional annotation. Nucleic Acids Res. 2016 Jan 4;44(D1):D733–45.

19. Hatcher EL, Zhdanov SA, Bao Y, Blinkova O, Nawrocki EP, Ostapchuck Y, et al. Virus Variation Resource – improved response to emergent viral outbreaks. Nucleic Acids Res. 2017 Jan 4;45(D1):D482–90.

20. Camacho CE, Boratyn G, Joukov V, Merezhuk Y, Madden T. ElasticBLAST [Internet]. NCBI; 2022. Available from: https://blast.ncbi.nlm.nih.gov/doc/elastic-blast/

21. Google Cloud Platform [Internet]. Google LLC; 2022. Available from: https://cloud.google.com/

22. Pace J, Youens-Clark K, Freeman C, Hurwitz B, Van Doorslaer K. PuMA: A papillomavirus genome annotation tool. Virus Evol. 2020 Jul 1;6(2):veaa068.

23. Hu JY, Hao ZQ, Frantz L, Wu SF, Chen W, Jiang YF, et al. Genomic consequences of population decline in critically endangered pangolins and their demographic histories. Natl Sci Rev. 2020 Apr 1;7(4):798–814.

24. Johnson M, Zaretskaya I, Raytselis Y, Merezhuk Y, McGinnis S, Madden TL. NCBI BLAST: a better web interface. Nucleic Acids Res. 2008 Jul 1;36(suppl_2):W5–9.

25. Shen W, Le S, Li Y, Hu F. SeqKit: a cross-platform and ultrafast toolkit for FASTA/Q file manipulation. PLOS ONE. 2016 Oct 5;11(10):e0163962.

26. Prjibelski A, Antipov D, Meleshko D, Lapidus A, Korobeynikov A. Using SPAdes de novo assembler. Curr Protoc Bioinforma. 2020;70(1):e102.

27. ORFfinder [Internet]. NCBI; 2022. Available from: https://www.ncbi.nlm.nih.gov/orffinder/

28. Gasteiger E, Gattiker A, Hoogland C, Ivanyi I, Appel RD, Bairoch A. ExPASy: the proteomics server for in-depth protein knowledge and analysis. Nucleic Acids Res. 2003 Jul 1;31(13):3784–8.

29. Boratyn GM, Thierry-Mieg J, Thierry-Mieg D, Busby B, Madden TL. Magic-BLAST, an accurate RNA-seq aligner for long and short reads. BMC Bioinformatics. 2019 Jul 25;20(1):405.

30. Danecek P, Bonfield JK, Liddle J, Marshall J, Ohan V, Pollard MO, et al. Twelve years of SAMtools and BCFtools. GigaScience. 2021 Feb 1;10(2):giab008.

31. Van Doorslaer K, Li Z, Xirasagar S, Maes P, Kaminsky D, Liou D, et al. The Papillomavirus Episteme: a major update to the papillomavirus sequence database. Nucleic Acids Res. 2017 Jan 4;45(D1):D499–506.

32. Liu P, Qiu Y, Xing C, Zhou JH, Yang WH, Wang Q, et al. Detection and genome characterization of two novel papillomaviruses and a novel polyomavirus in tree shrew (*Tupaia belangeri chinensis*) in China. Virol J. 2019 Mar 18;16(1):35.

33. Katoh K, Standley DM. MAFFT Multiple sequence alignment software version 7: improvements in performance and usability. Mol Biol Evol. 2013 Apr 1;30(4):772–80.

34. Darriba D, Posada D, Kozlov AM, Stamatakis A, Morel B, Flouri. ModelTest-NG: a new and scalable tool for the selection of DNA and protein evolutionary models. Mol Biol Evol. 2020 Jan; 37(1): 291–294.

35. Ronquist F, Teslenko M, van der Mark P, Ayres DL, Darling A, Höhna S, et al. MrBayes 3.2: Efficient Bayesian phylogenetic inference and model choice across a large model space. Syst Biol. 2012 May 1;61(3):539–42.

36. Kearse M, Moir R, Wilson A, Stones-Havas S, Cheung M, Sturrock S, et al. Geneious Basic: an integrated and extendable desktop software platform for the organization and analysis of sequence data. Bioinformatics. 2012 Jun 15;28(12):1647–9.

37. de Villiers EM, Fauquet C, Broker TR, Bernard HU, zur Hausen H. Classification of papillomaviruses. Virology. 2004 Jun 20;324(1):17–27.

38. Lauber C, Seitz S. Opportunities and challenges of Data-Driven Virus Discovery. Biomolecules. 2022 Aug;12(8):1073.

39. Aswad A, Katzourakis A. A novel viral lineage distantly related to herpesviruses discovered within fish genome sequence data. Virus Evol. 2017 Jul 1;3(2):vex016.

40. Zayed AA, Wainaina JM, Dominguez-Huerta G, Pelletier E, Guo J, Mohssen M, et al. Cryptic and abundant marine viruses at the evolutionary origins of Earth’s RNA virome. Science. 2022 Apr 8;376(6589):156–62.

41. Edgar RC, Taylor J, Lin V, Altman T, Barbera P, Meleshko D, et al. Petabase-scale sequence alignment catalyses viral discovery. Nature. 2022 Feb;602(7895):142–7.

42. Neri U, Wolf YI, Roux S, Camargo AP, Lee B, Kazlauskas D, et al. Expansion of the global RNA virome reveals diverse clades of bacteriophages. Cell. 2022 Sep 25;S0092-8674(22)01118-7.

43. Kawasaki J, Kojima S, Tomonaga K, Horie M. Hidden viral sequences in public sequencing data and warning for future emerging diseases. mBio. 2021 Aug 17;12(4):e01638–21.

