## Supplementary information for "Discovery of novel papillomaviruses in the critically-endangered Malayan and Chinese pangolins"

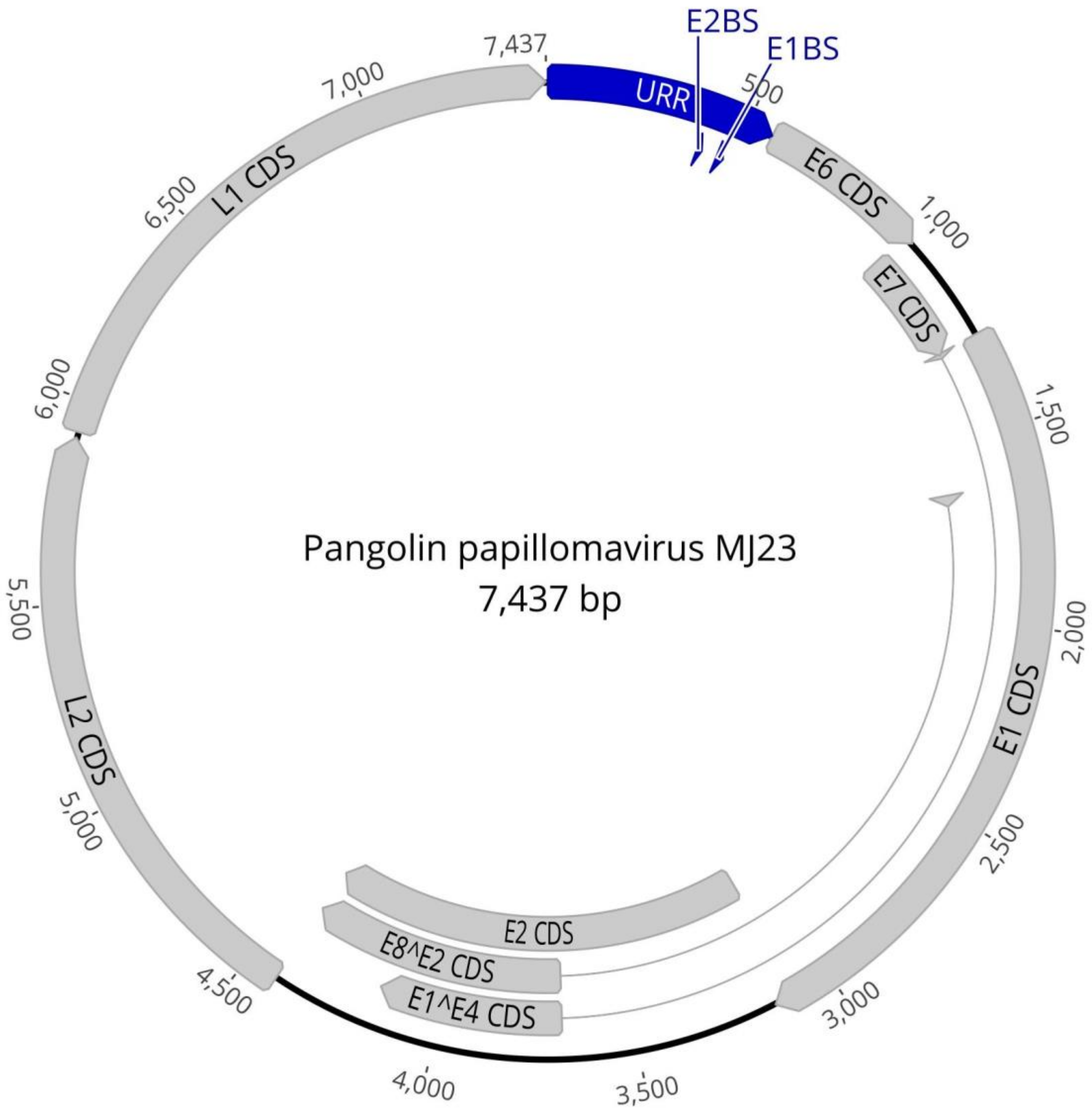


**Figure S1.** Complete papillomavirus genome assembled from the Malayan pangolin individual MJ23, and annotated with the PUMA pipeline. The 7,437-bp circular genome encodes the four core papillomavirus proteins (L1, L2, E1, E2) in addition to the E6 and E7 accessory proteins. Two spliced protein products were identified: E1^E4 and E8^E2. The Upstream Regulatory Region (URR) has also been annotated and includes the E1- and E2-protein binding sites (E1BS, E2BS). All genome features and sequences are provided as a GenBank file (Supplementary information). Image created in Geneious Prime 2022.2.1.

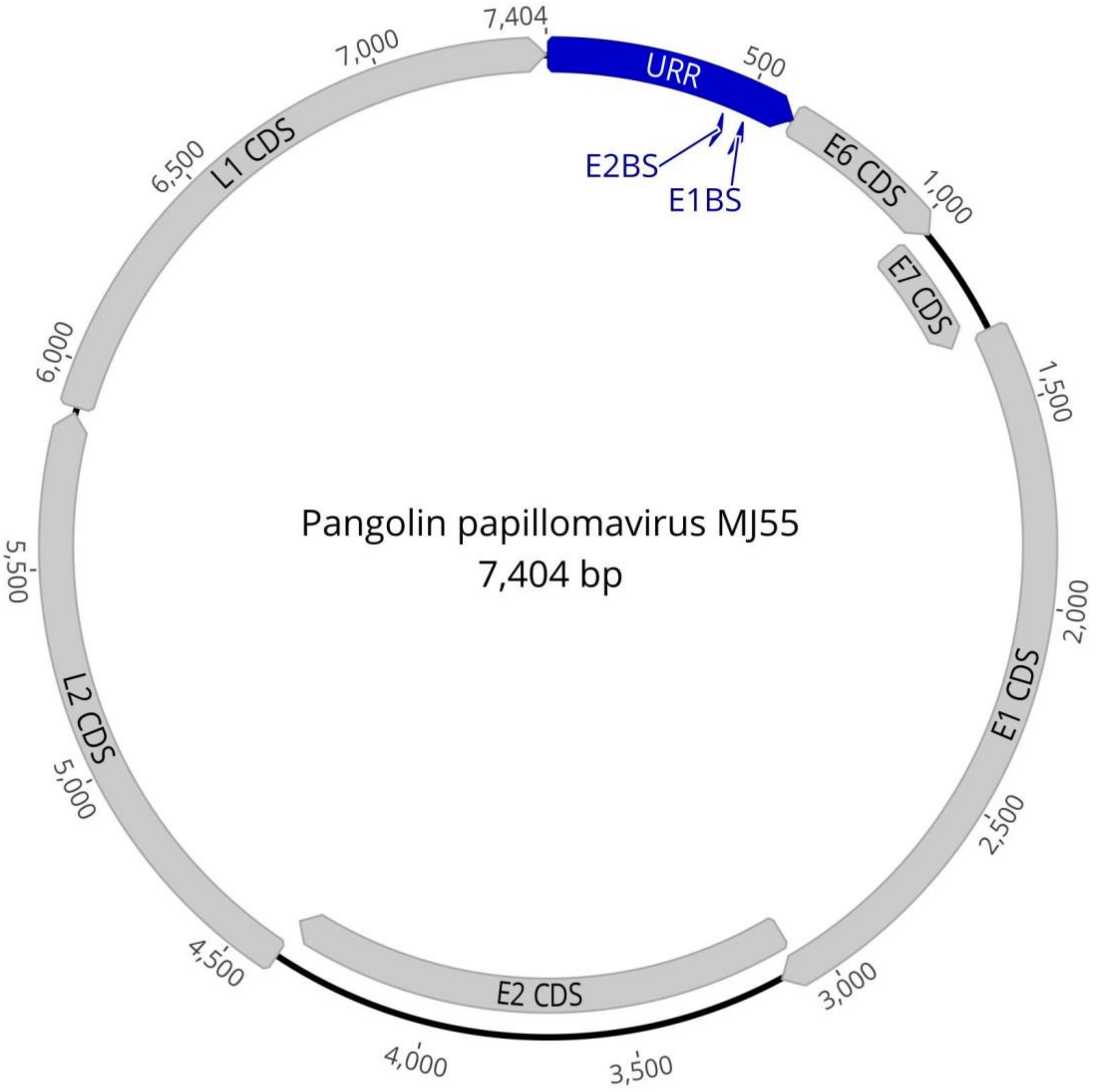


**Figure S2.** Papillomavirus genome assembled from the Malayan pangolin individual MJ23, and annotated with the PUMA pipeline. The 7,404-bp circular genome encodes the four core papillomavirus proteins (L1, L2, E1, E2) in addition to the E6 and E7 accessory proteins. Spliced protein products could not be identified. The Upstream Regulatory Region (URR) has also been annotated and includes the E1- and E2-protein binding sites (E1BS, E2BS). All genome features and sequences are provided as a GenBank file (Supplementary information). Image created in Geneious Prime 2022.2.1.

**Table S1.** Summary of the SRA experiments from pangolins with <70 significant hits (blastn, e-value < 0.01) to the linearised papillomavirus genome of the reference individual MJ74.

| Sample ID | Seizure location | Year | SRA accession | N. reads | % identity[1] |
|---|---|---|---|---|---|
| **MP14** | Yunnan, China | 2017 | SRR9018602 | 61 | 76.14 [69.50 - 87.25] |
| **MP03** | Yunnan, China* | 2000 | SRR9018585 | 50 | 76.21 [69.44 - 83.13] |
| **MJ61** | Yunnan, China | 2017 | SRR9018670 | 46 | 77.39 [69.17 - 100] |
| **MJ02** | Sino-Burmese border | 2014 | SRR9018651 | 42 | 82.61 [71.43 - 100] |
| **MJ03** | Sino-Burmese border | 2014 | SRR9018650 | 30 | 78.87 [70.91 - 100] |
| **MJ63** | Yunnan, China | 2016 | SRR9018672 | 23 | 76.47 [68.75 - 82.39] |
| **MJ41** | Sino-Burmese border | 2014 | SRR9018618 | 18 | 78.53 [71.90, 86.00] |
| **MJ04** | Sino-Burmese border | 2014 | SRR9018657 | 15 | 82.82 [71.55 - 100] |
| **MJ62** | Yunnan, China | 2016 | SRR9018673 | 12 | 82.57 [70.42 - 100] |
| **MP13** | Yunnan, China | 2016 | SRR9018605 | 11 | 75.56 [71.05 - 82.88] |
| **MJ51** | Yunnan, China | 2016 | SRR9018623 | 10 | 76.78 [72.00 - 85.81] |
| **MJ66** | Yunnan, China | 2017 | SRR9018587 | 9 | 76.98 [68.92 - 86.01] |
| **MP05** | Yunnan, China* | 2000 | SRR9018595 | 7 | 85.38 [76.98 - 100] |
| **MJ05** | Sino-Burmese border | 2014 | SRR9018656 | 7 | 75.84 [72.45 - 79.51] |
| **MP17** | Sino-Burmese border | 2014 | SRR9018609 | 6 | 84.87 [74.47 - 100] |
| **MJ73** | Yunnan, China | 2017 | SRR9018597 | 6 | 74.42 [71.71 - 76.74] |
| **MP18** | Sino-Burmese border | 2014 | SRR9018606 | 4 | 75.22 [73.64 - 76.60] |
| **MP16** | Sino-Burmese border | 2014 | SRR9018608 | 4 | 81.34 [78.05 - 84.62] |
| **MJ20** | Yunnan, China | 2017 | SRR9018635 | 4 | 74.63 [69.13 - 78.77] |
| **MJ28** | Sino-Burmese border | 2014 | SRR9018639 | 4 | 81.22 [77.42 - 89.13] |
| **MJ10** | Sino-Burmese border | 2014 | SRR9018666 | 4 | 78.98 [77.66 - 80.80] |
| **MP11** | Yunnan, China* | 2005 | SRR9018584 | 3 | 77.28 [74.29 - 79.17] |
| **MJ34** | Sino-Burmese border | 2014 | SRR9018642 | 3 | 74.41 [72.63 - 76.42] |
| **MJ01** | Sino-Burmese border | 2014 | SRR9018652 | 3 | 91.62 [80.46 - 97.60] |
| **MJ19** | Sino-Burmese border | 2014 | SRR9018661 | 3 | 75.49 [73.28 - 77.03] |
| **MJ16** | Yunnan, China* | 2000 | SRR9018664 | 3 | 80.01 [75.65 - 82.86] |
| **MP08** | Yunnan, China* | 2000 | SRR9018593 | 2 | 75.51 [75.51 - 75.51] |
| **MJ52** | Yunnan, China | 2016 | SRR9018624 | 2 | 78.07 [77.18 - 78.95] |
| **MJ21** | Sino-Burmese border | 2014 | SRR9018634 | 2 | 75.11 [71.96 - 78.26] |
| **MJ35** | Sino-Burmese border | 2014 | SRR9018643 | 2 | 81.97 [76.54 - 87.39] |
| **MJ42** | Sino-Burmese border | 2014 | SRR9018617 | 1 | 85.22 |
| **MJ22** | Sino-Burmese border | 2014 | SRR9018637 | 1 | 71.93 |
| **MJ32** | Sino-Burmese border | 2014 | SRR9018644 | 1 | 84.00 |

[1]Pairwise identity of the reads to the query sequence expressed as mean [min.-max.]
*Samples with known geographic origin.

**Table S2.** Characteristics of the open reading frames (ORFs) and predicted protein products encoded by the complete papillomavirus genomes and L1 *de novo* assemblies.

| Assembly | Contig length (bp) | Protein product | ORF Coordinates | Length (aa) | Molecular weight (Da) | Isoelectric point (pI) |
|---|---|---|---|---|---|---|
| **MJ74 complete genome[1]** | 7253 | E6 | 505–927 | 140 | 15799.42 | 8.72 |
| | | E7 | 929–1207 | 92 | 10292.84 | 4.05 |
| | | E1 | 1200–2999 | 599 | 69395.12 | 5.15 |
| | | E1^E4 | 1200–1212, 3537–3994 | 156 | 18747.84 | 9.64 |
| | | E8^E2 | 1576–1607, 3537–4239 | 244 | 26860.17 | 10.71 |
| | | E2 | 2941–4239 | 432 | 48840.44 | 9.40 |
| | | L2 | 4106–5722 | 538 | 59100.36 | 5.03 |
| | | L1 | 5739–7253 | 504 | 57268.48 | 6.13 |
| **MJ23 complete genome** | 7437 | E6 | 563–991 | 142 | 16275.00 | 9.07 |
| | | E7 | 967–1266 | 99 | 11103.94 | 4.13 |
| | | E1 | 1259–3148 | 629 | 72663.81 | 5.59 |
| | | E1^E4 | 1259–1274, 3686–4173 | 167 | 19290.32 | 6.26 |
| | | E8^E2 | 1638–1669, 3686–4418 | 254 | 28339.93 | 11.65 |
| | | E2 | 3090–4418 | 442 | 50569.74 | 9.92 |
| | | L2 | 4420–5907 | 495 | 53621.40 | 4.77 |
| | | L1 | 5923–7437 | 504 | 57635.88 | 6.23 |
| **MJ55 complete genome** | 7404 | E6 | 614–1042 | 142 | 16275.00 | 9.07 |
| | | E7 | 1018–1317 | 99 | 11103.94 | 4.13 |
| | | E1 | 1310–3121 | 603 | 69671.73 | 5.21 |
| | | E2 | 3063–4391 | 442 | 50613.75 | 9.91 |
| | | L2 | 4393–5880 | 495 | 53703.50 | 4.78 |
| | | L1 | 5896–7404 | 502 | 57312.53 | 5.99 |
| **MJ18** | 1577 | L1 | 1523–15* | 502 | 57028.24 | 6.30 |
| **MJ33** | 2094 | L1 | 292–1806 | 504 | 57635.88 | 6.23 |
| **MP15A** | 1664 | L1 | 121–1629 | 502 | 57028.24 | 6.30 |
| **MP15B** | 2059 | L1 | 238–1752 | 504 | 57635.88 | 6.23 |
| **MP15C** | 1727 | L1 | 1661–81* | 526 | 58702.52 | 7.50 |

[1]Reassembled genome for individual MJ74.
*ORF is on the reverse strand.

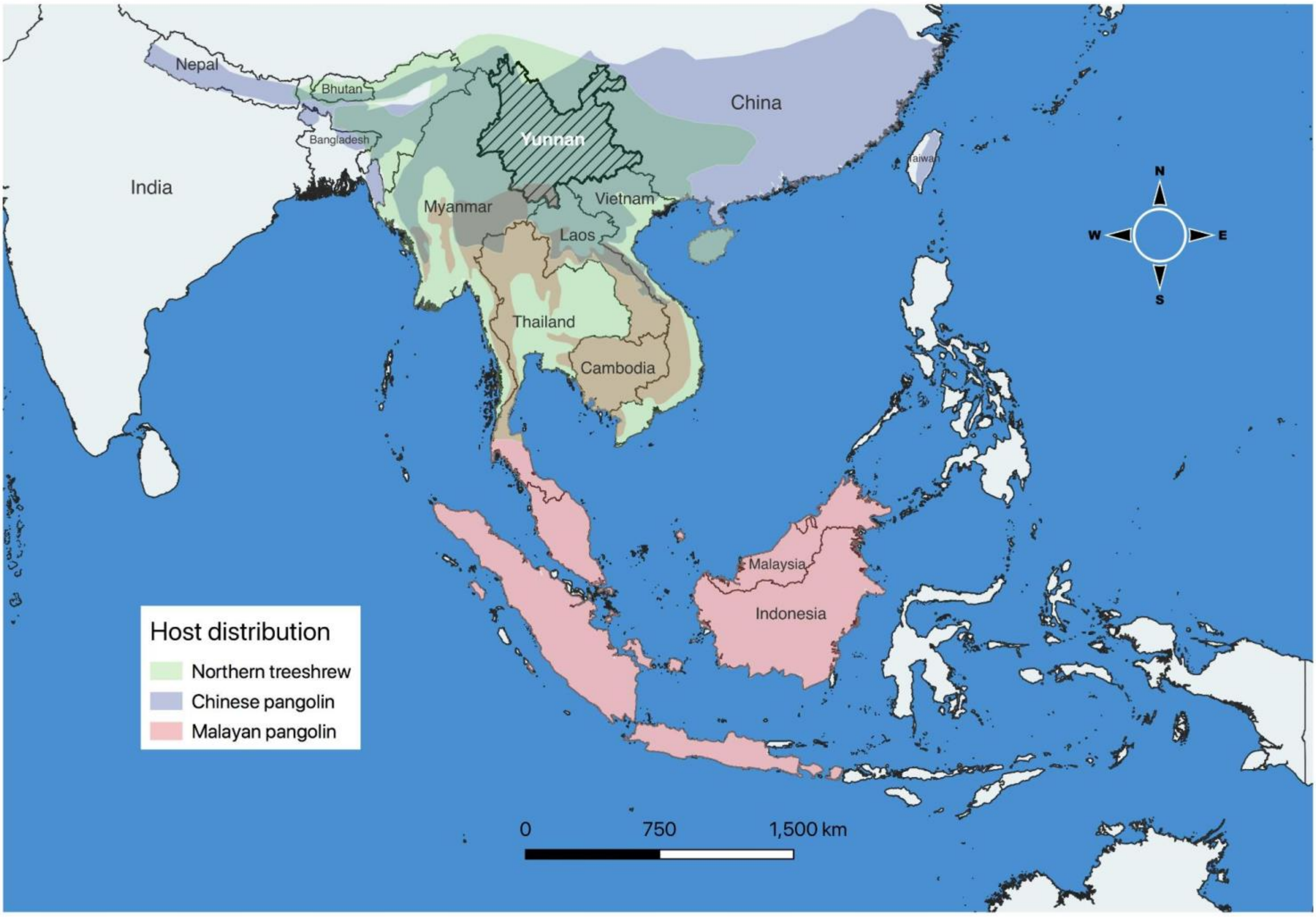


**Figure S3.** Map showing the ranges and overlap in the distributions of the Northern tree shrew (*Tupaia belangeri*), Chinese pangolin (*Manis pentadactyla*) and Malayan pangolin (*Manis javanica*) in South East Asia. The ranges of all three species overlap in regions of Myanmar, Northern Thailand, Laos, Central Vietnam and Yunnan, China. The Yunnan province of China is highlighted since all the pangolin individuals with positive reads to papillomaviruses and known geographic origin came from this region. All the other pangolin individuals with papillomavirus-positive reads were seized either in Yunnan or at the neighbouring border with Myanmar (Sino-Burmese border). Samples from individuals found in Guangzhou (4 *M. javanica*, seized), Malaysia (1 *M. javanica*, known location), Myanmar (1 *M. javanica*, known location) and Taiwan (1 *M. pentadactyla*, known location) were negative. Map produced in QGIS (1), with data from DIVA-GIS and the IUCN Red List (2–4).

**Table S3.** SRA experiments with significant hits to the MJ74 papillomavirus genome, found in the meta-transcriptomic study of the Malayan pangolin by Shi et al. (2022, doi: 10.1038/s41564-022-01181-1).

| Sample ID | Seizure location | Year | SRA accession | N. reads | % Identity[1] |
|---|---|---|---|---|---|
| **P162T** | Guangxi, China | 2018-2019 | SRR19632901 | 18 | 76.93 [71.81-83.09] |
| **P168T** | Guangxi, China | 2018-2019 | SRR19632895 | 2 | 75.48 [74.77-76.19] |
| **P097T** | Guangxi, China | 2018-2019 | SRR19632957 | 2 | 73.20 [70.75-75.64] |
| **P173T** | Guangxi, China | 2018-2019 | SRR19632890 | 1 | 72.39 |
| **P138T** | Guangxi, China | 2018-2019 | SRR19632925 | 1 | 76.60 |

[1]Pairwise identity of the reads to the query sequence expressed as mean [min.-max.]

**Table S4.** Best hits of the papillomavirus sequences reported in the publication of Liu et al. (2019, Table S1/S2, doi: 10.3390/v11110979), suggest those reads originate from human DNA contamination.

| GenBank accession | Virus | Description | Source | Notes |
|---|---|---|---|---|
| D00735.1 | HPV16 | Human papillomavirus type 16 genes for E6 protein, E7 protein, E1-E4 fusion protein, partial and complete cds | Vulvar intraepithelial neoplasia cell line (SK-v) | Contains human Alu element 802-1055 (AluJr subfamily) |
| D00486.1 | HPV16 | Human papillomavirus type 16 DNA, 5' virus-host integrated junction | Cervical carcinoma cell line (QG-U) | 189 bp 99% identical to gene *SLC36A1* (Chr5 5q33.1) |
| HE984519.1 | HPV16 | Human papillomavirus type 16 proviral 3' integration site, DNA junction sequence 2085_DJ2 | Cervical carcinoma | Human sequence flanking integrated HPV16 (Chr16 p13.13) |
| HE984542.1 | HPV16 | Human papillomavirus type 16 proviral 3' integration site, DNA junction sequence 3427_DJ1 | Cervical carcinoma | Human sequence flanking integrated HPV16 (Chr12 p13.2) |
| HE984550.1 | HPV16 | Human papillomavirus type 16 proviral 3' integration site, DNA junction sequence 4024_DJ2 | Cervical carcinoma | Human sequence flanking integrated HPV16 (Chr4 4q23) |
| HE984554.1 | HPV16 | Human papillomavirus type 16 proviral 3' integration site, DNA junction sequence 4749_DJ1 | Cervical carcinoma | Human sequence flanking integrated HPV16 (Chr18 p11.32) |
| HE984563.1 | HPV16 | Human papillomavirus type 16 proviral 3' integration site, DNA junction sequence 5189_DJ2 | Cervical carcinoma | Human sequence flanking integrated HPV16 (Chr8 q23.2) |
| HE984567.1 | HPV16 | Human papillomavirus type 16 proviral 3' integration site, DNA junction sequence CS_DJ2 | Cervical carcinoma cell line (CaSki) | Human sequence flanking integrated HPV16 (Chr10 p14) |
| X94741.1 | HPV16 | Human papillomavirus type 16 DNA for integration site of deleted allele | Squamous cell cervical carcinoma (SiHa) | DNA integration site of deleted allele (Chr13 q21-31) |

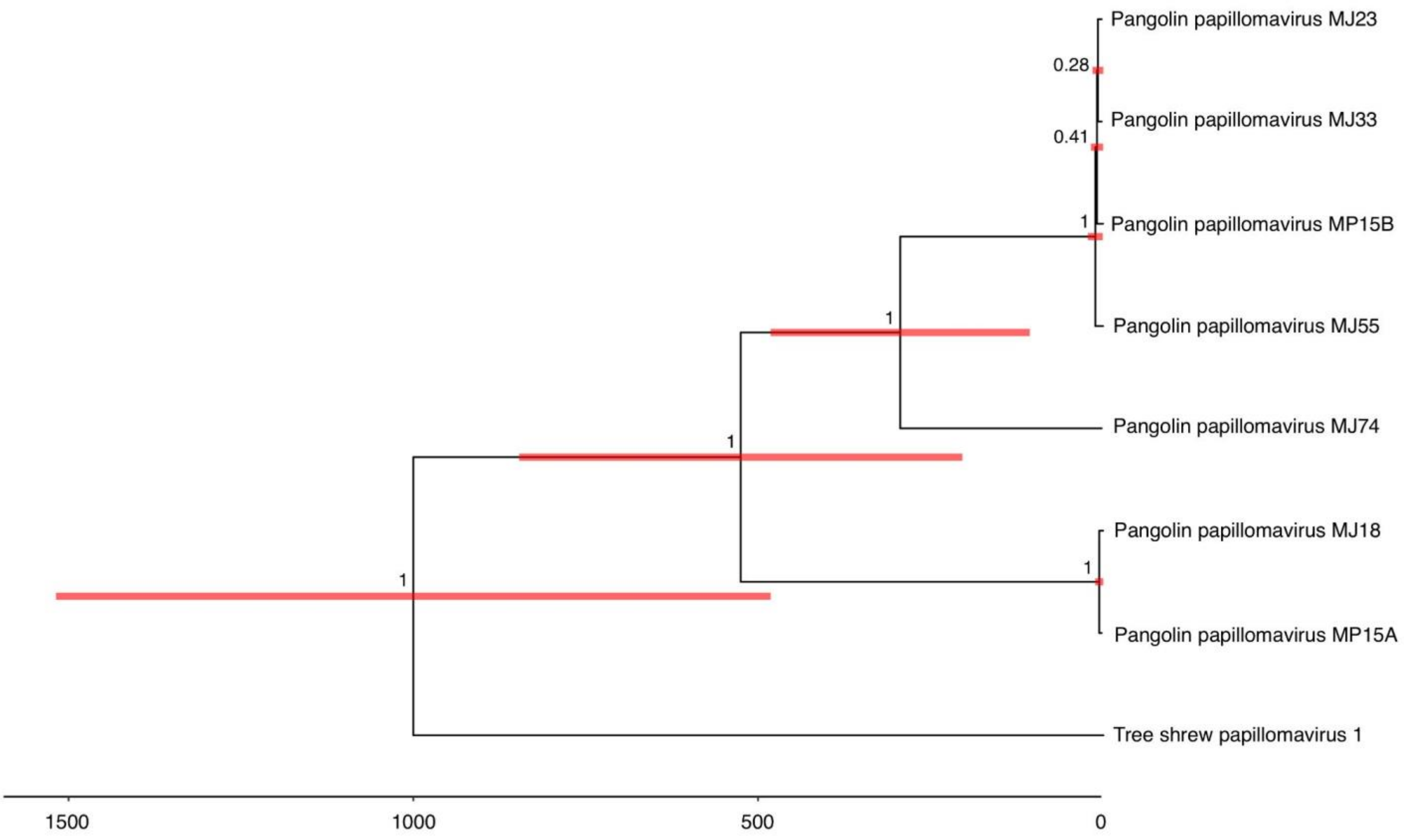


**Figure S4.** Bayesian maximum-credibility tree showing the relationships and divergence times of the tree shrew and pangolin papillomaviruses. The tree is based on an alignment of the 1st codon position of the L1 gene (513 characters), which showed a temporal signal using the sequence sampling times in TempEst ($R^2$ = 0.33) (5). The inference was carried out using a strict molecular clock, which was estimated as 3.03 (±0.96) × $10^{-4}$ nucleotide substitutions/site/year. The age of the root was estimated at 1001 (95% HPD: 638-1674) years, while the age of the most recent common ancestor of the pangolin papillomaviruses was estimated at 526 (95% HPD: 262-905) years. Tree estimated in BEAST2 (6). MCMC chain length = 20 million generations, burn-in = 25%.

**Supplementary text**

Data collection: the raw data analysed in this study is accessible under the BioProject identifiers PRJNA529512, PRJNA529540, PRJNA845961 and PRJNA573298 on the NCBI website (https://www.ncbi.nlm.nih.gov/).

File formats used to record the data: short-reads were downloaded in *.fasta format. The calculation of mapping reads and coverage used files in the *.sam and *.bam formats. The multiple sequence alignment used in MrBayes was in *.nexus format. The input file for the analysis of the timetree in BEAST is in *.xml format. The consensus and maximum credibility Bayesian trees are in *newick format.

Description of columns and tables in data set: column definitions for the *.sam files are as follows: column 1 = sequence id, column 2 = bitwise FLAG, column 3 = reference id, column 4 = leftmost mapping position, column 5 = mapping quality, column 6 = CIGAR string, column 7 = name of mate/next read, column 8 = position of mate/next read, column 9 = observed template length, column 10 = segment sequence, column 11 = ASCII of Phred score (https://samtools.github.io/hts-specs/SAMv1.pdf).

Details of scripts and code: the python script used to parse paired-end reads into forward, reverse and orphan files is available on Github (https://github.com/josegabrielnb/pangolins).

Details of software used:

ElasticBLAST v0.2.6 (https://blast.ncbi.nlm.nih.gov/doc/elastic-blast/)
Google Cloud Platform (https://cloud.google.com)
PuMA annotation pipeline v1.2.1 (https://github.com/KVD-lab/puma)
BLASTN/TBLASTN hosted on the NCBI web server (https://blast.ncbi.nlm.nih.gov/Blast.cgi)

SeqKit v2.2.0 (https://bioinf.shenwei.me/seqkit/)

Python v3.7.9 (https://www.python.org/downloads/)

SPAdes v3.15.4 (https://github.com/ablab/spades)

ORFfinder (https://www.ncbi.nlm.nih.gov/orffinder/)

ProtParam, Expasy (https://web.expasy.org/protparam/)

MagicBLAST v1.6.0 (https://ncbi.github.io/magicblast/)

Samtools v1.15.1 (https://github.com/samtools/samtools)

Papillomavirus Episteme, PaVE (https://pave.niaid.nih.gov/)

MAFFT v7.490 (https://mafft.cbrc.jp/alignment/software/)

ModelTest-NG v0.1.7 (https://github.com/ddarriba/modeltest)

MrBayes v3.2.7a (https://nbisweden.github.io/MrBayes/)

TempEst v1.5.3 (http://tree.bio.ed.ac.uk/software/tempest/)

BEAST2 v2.6.7 (https://www.beast2.org/)

Tracer v1.7.2 (https://github.com/beast-dev/tracer/releases/tag/v1.7.2)

QGIS v3.26 Buenos Aires (https://www.qgis.org/en/site/)

Description of how data was processed to obtain results + statistical tests: reads with significant matches to our papillomavirus queries were downloaded as *.fasta files. A custom python script was used to sort the reads into forward, reverse and orphan files (see Details of script and code). These files were used as the input for assembly in SPAdes with the "-1", "-2" and "-s" flags, respectively. For the calculation of the depth of assembled sequences, the reads were mapped to the reference assembly for that individual, and the results were stored as *.sam files. These files were converted to the *.bam format for sorting the reads and then calculating the depth. The MAFFT multiple sequence alignment was trimmed manually to remove poorly aligned columns, resulting in a 522 aa alignment for the L1 protein. The trimmed alignment was converted into *.nexus format for use in MrBayes. Convergence in BEAST2 was confirmed by examination of the MCMC on Tracer, ensuring good mixing, stationarity and that the effective sample sizes (ESSs) for all parameters > 200.

Licences/restrictions placed on data sets: all the raw sequencing data sets are freely accessible online. The data associated with our analysis will be made freely available in Dryad and the assembled sequences will be deposited in GenBank.

**Supplementary material references**


1. QGIS Development Team. QGIS Geographic Information System. Open Source Geospatial Foundation Project. [Internet]. Available from: http://qgis.osgeo.org
2. Challender D, Willcox DHA, Panjang E, Lim N, Nash H, Heinrich S, et al. *Manis javanica*. The IUCN Red List of Threatened Species 2019: e.T12763A123584856. [Internet]. Available from: https://dx.doi.org/10.2305/IUCN.UK.2019-3.RLTS.T12763A123584856.en
3. Challender D, Wu S, Kaspal P, Khatiwada A, Ghose A, Ching-Min S, et al. *Manis pentadactyla* (errata version published in 2020). The IUCN Red List of Threatened Species 2019: e.T12764A168392151. [Internet]. Available from: https://dx.doi.org/10.2305/IUCN.UK.2019-3.RLTS.T12764A168392151.en
4. Han KH, Duckworth JW, Molur S. *Tupaia belangeri*. The IUCN Red List of Threatened Species 2016: e.T41492A22280884 [Internet]. 2016 [cited 2022 Aug 18]. Available from: https://dx.doi.org/10.2305/IUCN.UK.2016-2.RLTS.T41492A22280884.en
5. Rambaut A, Lam TT, Max Carvalho L, Pybus OG. Exploring the temporal structure of heterochronous sequences using TempEst (formerly Path-O-Gen). Virus Evol. 2016 Jan 1;2(1):vew007.
6. Bouckaert R, Heled J, Kühnert D, Vaughan T, Wu C-H, Xie D, et al. BEAST 2: a software platform for Bayesian evolutionary analysis. PLoS Comput Biol. 2014 Apr 10; 10(4): e1003537.
