## Supplementary material for "Discovery of novel papillomaviruses in the critically-endangered Malayan and Chinese pangolins": sort_reads.py

```
#!/usr/local/bin/python3

#Read a fasta file of short reads and sort them into forward, reverse and orphan fasta files

from Bio import SeqIO

prelim_forward = []
prelim_reverse = []

for fasta in SeqIO.parse(open("all_reads-noduplicates.fasta"),'fasta'):

    if fasta.id.endswith(".1"):

        prelim_forward.append(fasta.id[:-1])

    elif fasta.id.endswith(".2"):

        prelim_reverse.append(fasta.id[:-1])

#print(prelim_forward)
#print(prelim_reverse)

added = []

for fasta in SeqIO.parse(open("all_reads-noduplicates.fasta"),'fasta'):
    
    if fasta.id[:-1] in prelim_forward and fasta.id[:-1] in prelim_reverse\
         and fasta.id not in added:

        if fasta.id.endswith(".1"):

            with open("forward_reads.fasta","a") as f:
                f.write(">"+fasta.id+"\n")
                f.write(str(fasta.seq)+"\n")
            f.close()

        elif fasta.id.endswith(".2"):

            with open("reverse_reads.fasta","a") as f:
                f.write(">"+fasta.id+"\n")
                f.write(str(fasta.seq)+"\n")
            f.close()

        added.append(fasta.id)

    else:

        with open("orphan_reads.fasta","a") as f:
                f.write(">"+fasta.id+"\n")
                f.write(str(fasta.seq)+"\n")
        f.close()
```
