## Supplementary material for "Discovery of novel papillomaviruses in the critically-endangered Malayan and Chinese pangolins": README.txt

The .fsa and .tbl files in this folder are for the genbank submission process.
Please refer to https://www.ncbi.nlm.nih.gov/genbank/tbl2asn2/ to utilize the files created with PuMA along with user created files to start the genbank submission process.
